# Molecular dynamics simulations reveal membrane lipid interactions of the full-length lymphocyte specific kinase Lck

**DOI:** 10.1101/2022.05.10.491278

**Authors:** Dheeraj Prakaash, Graham P. Cook, Oreste Acuto, Antreas C. Kalli

## Abstract

The membrane-bound lymphocyte-specific protein-tyrosine kinase (Lck) triggers T cell antigen receptor signalling to initiate adaptive immune responses. Despite many structure-function studies, the mode of action of Lck and the potential role of plasma membrane lipids in regulating Lck’s activity remains elusive. Advances in molecular dynamics simulations of membrane proteins in complex lipid bilayers have opened a new perspective in gathering such information. Here, we have modelled the full-length Lck open and closed conformations available from crystallographic studies and simulated its interaction with the inner leaflet of the T cell plasma membrane. In both conformations, we found that the unstructured unique domain and the structured domains including the kinase interacted with the membrane with a preference for PIP lipids. Interestingly, our simulations suggest that the Lck-SH2 domain interacts with lipids differently in the open and closed Lck conformations, demonstrating that lipid interaction can potentially regulate Lck’s conformation and in turn modulate T cell signalling. Additionally, the Lck-SH2 and kinase domain residues that significantly contacted PIP lipids are found to be conserved among the Src family of kinases, thereby potentially representing similar PIP interactions within the family.

## INTRODUCTION

Activation of T cells is triggered by the engagement of the T cell receptor (TCR) with antigenic peptides presented by major histocompatibility complexes (pMHC) (Courtney, Lo, & Weiss, 2018; Mariuzza, Agnihotri, & Orban, 2019). Upon pMHC binding, allosteric sites in the extracellular and transmembrane regions of the T cell receptor-CD3 complex (TCR-CD3) (He et al., 2020; Lanz et al., 2021) promote exposure of immunoreceptor tyrosine-based activation motifs (ITAMs) in the cytoplasmic tails of CD3/ζ subunits. Further, ITAMs are promptly phosphorylated by Lck. Remarkably, non-activated T cells maintain a sizable fraction of constitutively activated Lck at the plasma membrane that is necessary and sufficient for ITAM phosphorylation upon ligand binding (Nika et al., 2010). Additionally, imaging studies have suggested that Lck conformational states dictate its spatial distribution that may impact TCR-CD3 ITAM phosphorylation (Rossy, Owen, Williamson, Yang, & Gaus, 2013). Phosphorylated ITAMs then provide stable binding sites for the tyrosine kinase ZAP-70 (Hatada et al., 1995; Katz, Novotná, Blount, & Lillemeier, 2017) that is regulated by Lck to propagate signals required for T cell activation (Palacios & Weiss, 2004).

Understanding the role of Lck in molecular detail is important in deciphering the initial phases of T cell activation. To achieve this, it is key to obtain the full-length 3D structure of Lck which remains structurally unresolved until date. The full-length Lck contains the following domains (from the *N* to *C* terminus): the SH4 (first ~10 residues), unique domain (UD; following ~50 residues) both of which are likely to be devoid of secondary structure. The UD is followed by the SH3, SH2, and the kinase domains for which structural data is available. The X-ray crystallographic structure of the Lck-SH2 and SH3 domains combined is available at a resolution of 2.36 Å (PDB:4D8K), and the isolated kinase domain in its active state i.e., phosphorylated at Y394 is available at 1.7 Å resolution (PDB:3LCK) (Yamaguchi & Hendrickson, 1996). However, the structures of the SH4 and UD remain largely unresolved. Nonetheless, NMR data for Lck-UD in solution indicates that it lacks structure and has no significant influence on Lck-SH3 (Briese & Willbold, 2003). In this work, we refer to the SH4 and the UD combined as the ‘SH4-U’ domain for simplicity. In the SH4 domain, G2, C3 and C5 undergo acylation as a post-translational acylation i.e., myristoylation at G2 (Udenwobele et al., 2017; Wingfield, 2017), and palmitoylation at C3 and C5 (Yurchak & Sefton, 1995). As a result, the acyl chains or lipid tails covalently attached to these residues insert into the hydrophobic core of the membrane and aid in membrane localization of Lck (MD, 1994).

Lipids in the inner leaflet of the plasma membrane have been reported to play an important role in interacting with Lck via its SH2 domain and in turn regulating TCR-CD3 signalling (Sheng et al., 2016). In particular, anionic lipids such as phosphatidylinositol-4,5-biphosphate (PI(4,5)P2 or PIP_2_) and phosphatidylinositol-3,4,5-triphosphate (PIP_3_) were suggested to aid in Lck interaction with the TCR-CD3 in a spatiotemporal manner. Lck-SH2 domain is found to be key to lipid interaction by selectively contacting these PIP lipids via a cationic patch (Sheng et al., 2016). SH2 domains in other tyrosine kinases such as Zap70 have also been reported to direct signalling pathways by binding to PIP lipids (Park et al., 2016). Our previous studies suggested that the TCR-CD3 maintains an anionic lipid environment enriched in PIP lipids with the help of its cytoplasmic region (Prakaash, Cook, Acuto, & Kalli, 2021). Since Lck is also shown to possess high affinity for PIP lipids (Sheng et al., 2016) and its clustering is driven by their open conformational state (Rossy et al., 2013), it is important to understand how the open and closed states of Lck interact with the membrane in molecular detail. This could further aid in our understanding of its interaction with ITAMs of stimulated TCR-CD3 complexes.

In this study, we modelled the full-length Lck by predicting the structure of the SH4-U domain and integrating it with the experimentally resolved structures of the SH2, SH3 and kinase domains. Further, we performed coarse-grained molecular dynamics simulations over a cumulative time of 100 microseconds for each of the open and closed states of Lck in a complex symmetric bilayer whose lipid headgroup composition resembles the inner leaflet of the T cell plasma membrane (Zech et al., 2009). From these simulations, we study the conformational dynamics and lipid interactions of the open and closed conformations of the full-length Lck.

## RESULTS and DISCUSSION

### Modelling the full-length Lck in its open and closed states

To obtain a model of the 3D structure of full-length Lck, we first modelled the SH4-U domain since its structure is unknown. To achieve this, we used two independent 3D structure prediction tools i.e., I-Tasser (Yang et al., 2015) and Robetta (D. E. Kim, Chivian, & Baker, 2004), and obtained multiple 3D models from each. Then, the PSIPRED Protein Analysis Workbench (Buchan & Jones, 2019) was used to calculate the secondary structure of the SH4-U domain (**Fig S1A**). The best 3D structural models, one from I-Tasser and one from Robetta (representing the highest prediction confidence score and agreement with secondary structure predictions) were subjected to 250 ns atomistic molecular dynamics (ATMD) simulations in solution neutralized by 0.15M Na+ and Cl- ions to allow optimization of the predicted structures. At the end of the ATMD simulations, we used two criteria to select the best model: (i) agreement with secondary structure predictions, and (ii) agreement with structural information revealed by an NMR study (P. W. Kim, Sun, Blacklow, Wagner, & Eck, 2003) where the UD contained a hairpin-like loop region (**Fig 1A left**). This loop region was found to be responsible for binding CD4 and CD8 co-receptors via a coordinating Zn^2+^ ion (P. W. Kim et al., 2003), though data have indicated that not all Lck are bound to co-receptors. Using these criteria, the model derived from the Robetta server was selected. However, due to the absence of a zinc ion in our model, a disulphide bond was formed. In addition, our model suggests that residues E10, D11, D12, E15, E21 (**Fig 1A**) form an anionic patch that potentially interact with cationic regions of the TCR-CD3 cytoplasmic region. Moreover, the D12N mutation was shown to reduce binding with CD3ε BRS by NMR experiments (Li et al., 2017).

**Fig 1.**
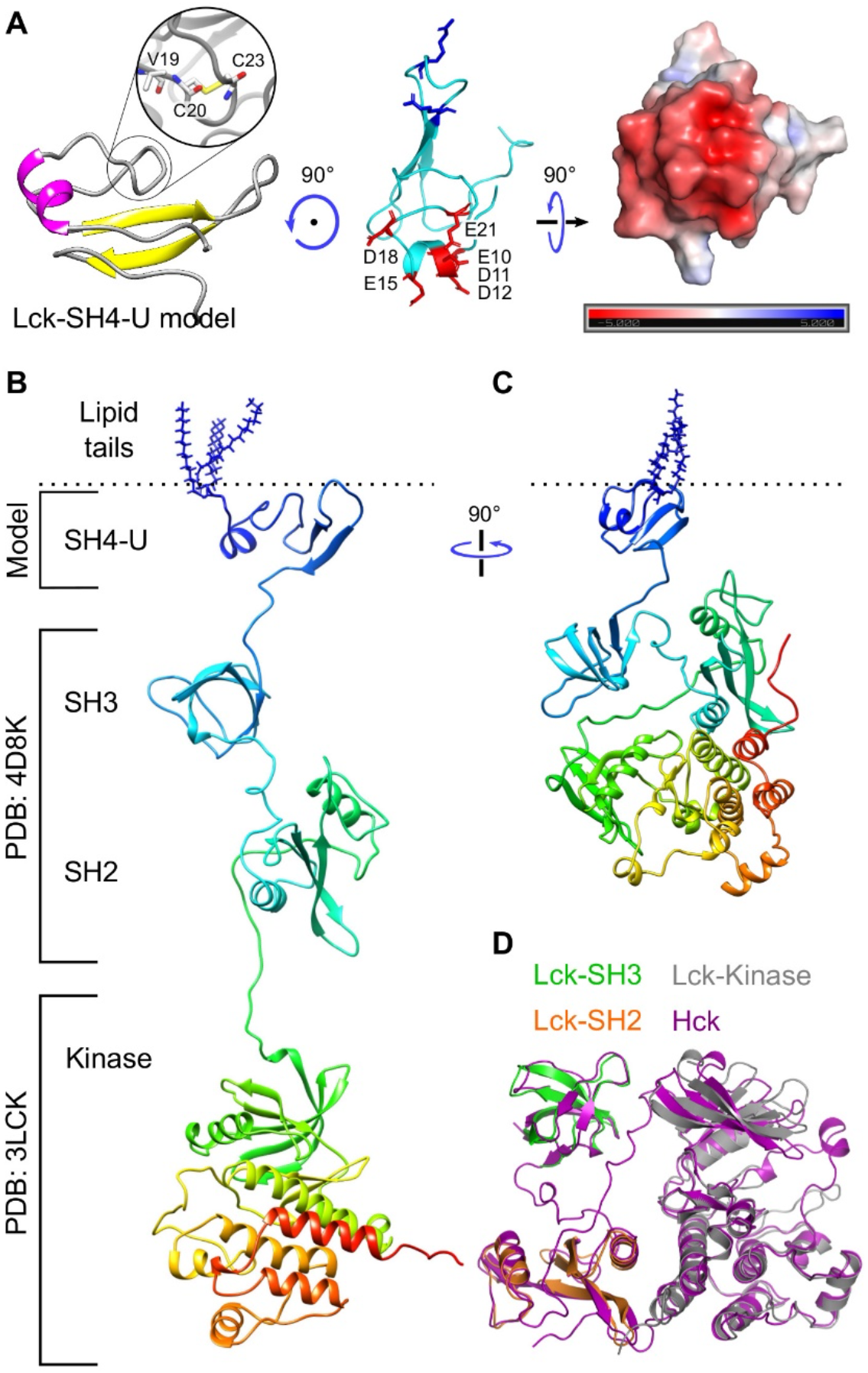
Model of the SH4-U domain and, the open and closed full-length Lck conformations. ***(A)** The model of the SH4-U domain (residues 2 to 63) used in this study. The region responsible for coordinating a Zn^2+^ ion is magnified (left). Residues forming the anionic patch in our model are shown as red sticks and labelled. PIP lipid binding cationic residues (R39, R45), as suggested in this study, are shown as blue sticks are located on the opposite side of the anionic patch (middle). The electrostatic profile of the anionic patch (right) shown was calculated in the ±5 kT/e range and at pH 7.0 using the PDB2PQR* (Dolinsky, Nielsen, McCammon, & Baker, 2004) *and APBS* (Baker, Sept, Joseph, Holst, & McCammon, 2001) *tools. Electronegative and electropositive regions are indicated by red and blue intensities respectively. **(B)** The model of the open Lck-FL conformation and **(C)** the closed Lck-FL conformation used in this study. **(D)** Isolated SH2, SH3 and kinase domains aligned to the closed state of Hck (PDB:5H0B)*.

Following modelling of the SH4-U domain, the crystal structures of SH2, SH3 (PDB:4D8K), and kinase domains (PDB:3LCK) along with the SH4-U model were assembled in a linear manner to model the full-length Lck open conformation (**Fig 1B**) using UCSF Chimera (Pettersen et al., 2004). Missing residues located in the linker region were predicted to be unstructured and hence modelled as loops between the domains using Modeller 9.2 (Webb & Sali, 2014). The structures of the different domains were assembled sufficiently far from each other to avoid bias in protein-protein interactions at the beginning of the simulations. Note that the positioning of the SH3 and SH2 domains relative to each other were not altered and were used as obtained from the crystal structure (PDB:4D8K).

The crystal structure of the closed conformation of Hck, an Src family member of kinases, resolved at 1.65 Å (PDB:5H0B) was used as a template to model the Lck closed state using Modeller 9.2 (Webb & Sali, 2014). The resultant homology model of the Lck closed state (containing SH2, SH3, kinase domains) was then conjoined with the SH4-U model (as shown in **Fig 1A**) to obtain the full-length Lck (Lck-FL) in its closed state (**Fig 1C**). This modelling used a multiple sequence alignment of Hck and Lck produced by Clustal Omega (Sievers & Higgins, 2018). The structures of the SH2, SH3, kinase domains of Lck were also individually aligned with those respective domains of Hck (**Fig 1D**) using the ‘super’ aligning method in PyMOL (pymol.org) indicating their structural similarities i.e., RMSD = 0.582, 1.135, 0.92 Å respectively.

Finally, to both the Lck-FL open and closed models, post-translational modifications were added to the *N*-terminal residues i.e., G2 was myristoylated and, C3 and C5 were palmitoylated prior to the simulations. The initiator Met1 residue was removed during this process since they are known to be cleaved in mature eukaryotic proteins (16, 17).

### Membrane association and lipid interaction of the full-length Lck

To assess the association of the Lck-FL models with the membrane (see **Table 1** for membrane composition), we performed coarse-grained molecular dynamics (CGMD) simulations. At the beginning of these simulations, the post-translational modifications (lipid tails) of both the Lck-FL open and closed models were made to partially penetrate the membrane surface to mimic the fact that the lipid tails are expected to penetrate the membrane upon binding of Lck to the membrane. 20 individual simulations for 5 μs were performed for both the open and closed models. Calculation of the average distance versus time of the center of mass (COM) of the initial protein model to the COM of the membrane along the vertical (Z) axis showed that both Lck-FL models closely associated with the membrane within 1 μs simulation time (**Fig 2A**). Analysis of Lck interactions with lipids shows an increase in the number of PIP_2_ and PIP_3_ around the protein in both its open and closed conformations creating an anionic annulus around the protein. This annulus was retained for the remaining time of the simulations (**Fig 2B**). For both models, radial distribution function (RDF) showed that PIP_2_ and PIP_3_ were preferred by Lck over other lipids (**Fig 2C**). Experimental studies also suggest that the SH2 domain prefers to interact with PIP_3_ compared to PI(4,5)P2 (Sheng et al., 2016). Interestingly, these lipid species were found to also cluster around the TCR-CD3 and interact with its cytoplasmic region as observed in our previous studies of the complete TCR-CD3 complex (Prakaash et al., 2021).

**Table 1.** Summary of CGMD simulations conducted in this study.

| Simulations | Membrane | Particles | Simulation box<br>(X × Y × Z axis) | Duration | Replicas |
| --- | --- | --- | --- | --- | --- |
| Lck-SH4-U | simple | 22134 | 12 × 12 × 17 | 1 μs | 20 |
| Lck-SH3 | simple | 25676 | 12 × 12 × 20 | 1 μs | 20 |
| Lck-SH2 | simple | 25914 | 12 × 12 × 20 | 1 μs | 20 |
| Lck-SH2,3,4-U | complex | 53576 | 16 × 16 × 23 | 5 μs | 20 |
| Lck-FL open | complex | 83137 | 19 × 19 × 26 | 5 μs | 20 |
| Lck-FL open mut5 | complex | 83194 | 19 × 19 × 26 | 5 μs | 20 |
| Lck-FL closed | complex | 46187 | 16 × 16 × 20 | 5 μs | 20 |

(Symmetric) membrane composition:
- simple: POPC/POPS/PIP<sub>2</sub>/PIP<sub>3</sub> = 72/20/6/2 - complex: POPC/POPE/POPS/Chol/PIP<sub>2</sub>/PIP<sub>3</sub> = 12/40/20/20/6/2

**Fig 2.**
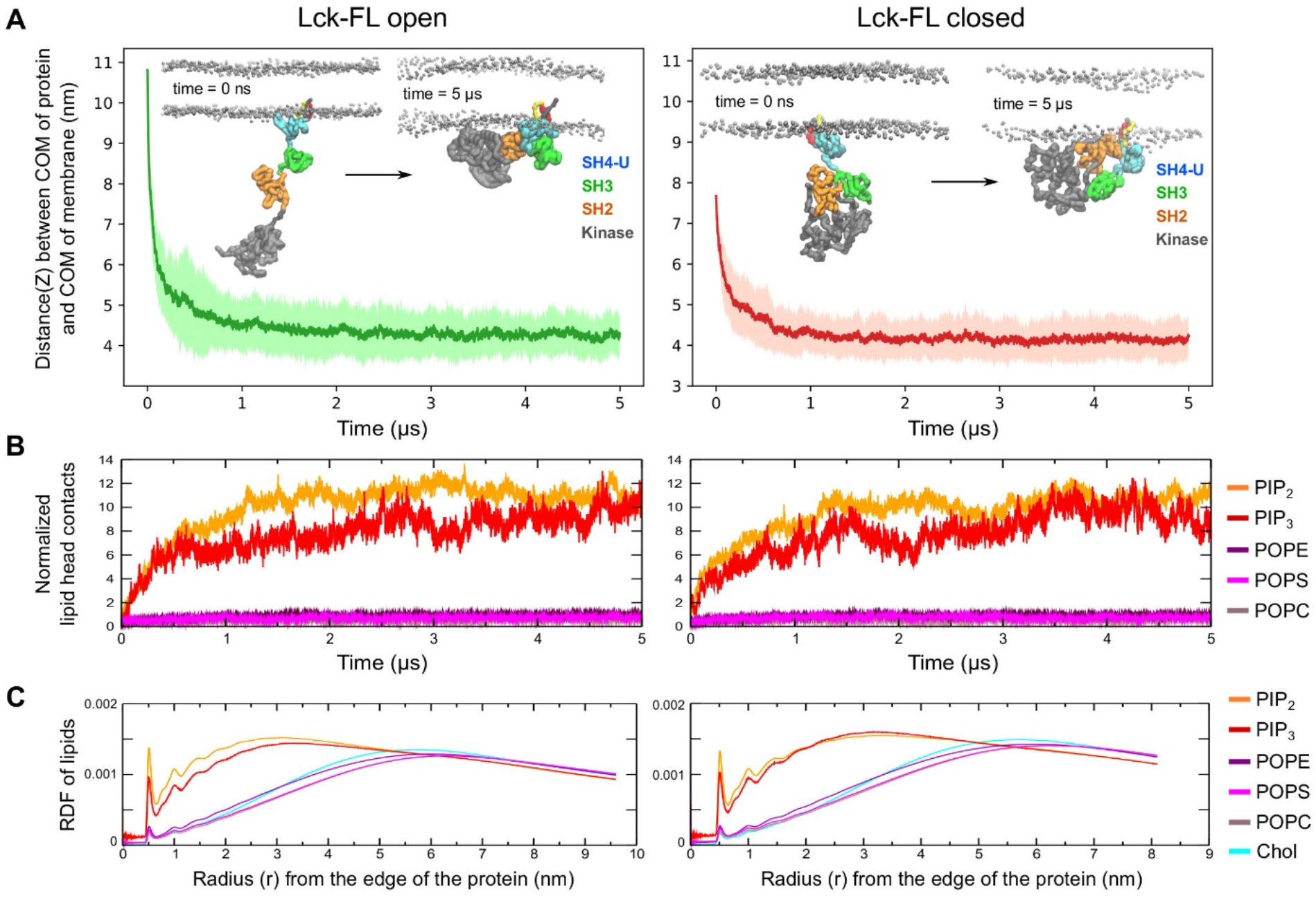
Membrane association and lipid interactions of the open and closed full-length Lck conformations. **(A)** Association of open and closed conformations of Lck-FL with the membrane is indicated by the reduction in distance between the center of mass (COM) of Lck-FL and COM of the membrane versus time. **(B)** Number of interactions between Lck and lipid headgroups versus time. The number of headgroup interactions of each phospholipid type is normalized by the number of lipids of the respective lipid type in the membrane. **(C)** The radial distribution function (RDF) of all lipid types around Lck calculated throughout the simulation time. The RDF is normalized by the total number of lipids in the membrane of that system to enable comparison between the open and closed conformations of Lck-FL. Note: Simulations of the Lck-FL open and closed systems contain different number of lipids in the membrane due to different sizes of the membrane.

Given the potential of strong electrostatic interactions between Lck and PIP lipids, and also between the TCR-CD3 cytoplasmic tails and PIP lipids of the inner leaflet of the membrane (Prakaash et al., 2021), it is possible that TCR-Lck association and ITAM phosphorylation occurs proximal to the inner leaflet of the plasma membrane.

### PIP lipid binding sites

Analysis of the interactions of the Lck-FL open conformation (initial simulation frame shown in **Fig 3A left**) with PIP lipids showed that the SH2 domain made significant contacts via a primary binding site (K182 > R184 ~ R134 > K135 ~ K179). The SH4-U domain also made significant contacts by preferring to bind to PIP lipids via R39 and R45, followed by cholesterol interactions via myristoylated G2, palmitoylated C3 and C5, followed by residues H24, Y25, P26, V44, R45, D46 **(Fig 3B)**. The SH3 domain made less contact with the membrane, interacting mostly via K84. Interestingly, the kinase domain of Lck also showed significant PIP lipid interactions; the most interactive residues were R455, R458, R474, K478 (**Fig 3B**). These residues are situated at the bottom surface of the C-terminal lobe of the kinase domain and constitute a flat cationic area acting as a PIP lipid binding site (**Fig S1B**). Furthermore, in our simulations, residues including and neighbouring A160 were observed to interact with lipids, but not as significantly as K182 and R184 (**Fig 3B**). This observation is consistent with mutation studies which revealed that A160K reduces dissociation of Lck-SH2 from plasma membrane-mimetic vesicles (Sheng et al., 2016).

**Fig 3.**
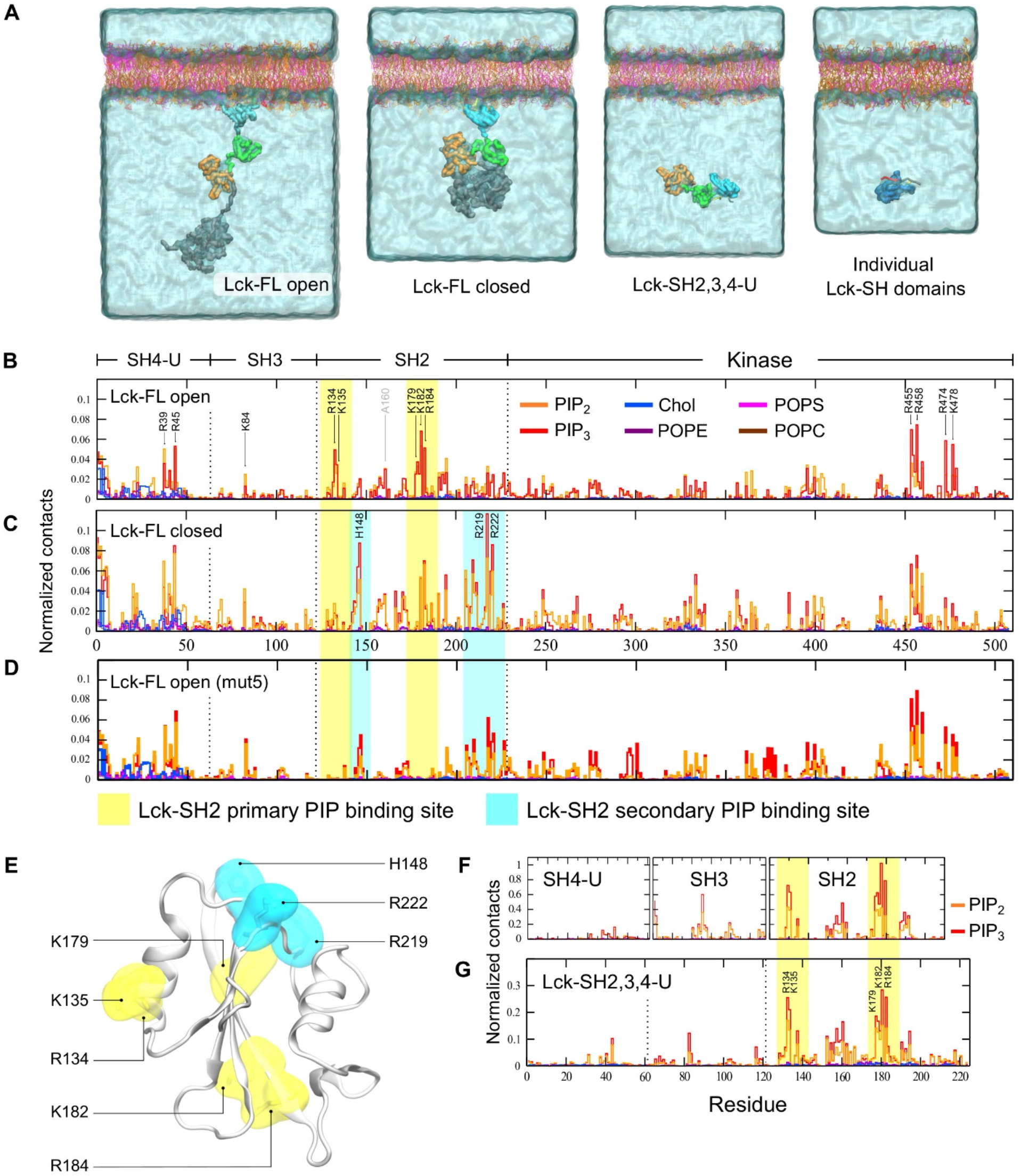
Snapshots of the initial simulation setup and PIP lipid binding sites. **(A)** The initial frames of the simulations of the open and closed conformations of Lck-FL showing the lipid tails of the SH4 domain partially inserted into the membrane at the beginning of simulation. Also, the initial frames of the simulations of the Lck-SH domains combined (SH2, SH3, SH4-U) and of the individual Lck-SH domains where the protein structure is placed in solution ~6 nm away from the membrane. **(B)** Normalized lipid interactions of Lck-FL open, **(C)** Lck-FL closed, **(D)** Lck-FL open when mutated (mut5 i.e., R134A, K135A, K179A, K182A, R184A), **(E)** Residues constituting the primary (yellow) and secondary (cyan) PIP lipid binding sites of Lck-SH2 as observed in CGMD simulations of Lck-FL open and closed. **(F)** Normalized lipid interactions of the SH4-U, SH3, SH2 domains when individually simulated, and **(G)** of the SH4-U, SH3, SH2 domains simulated in conjunction (Lck-SH2,3,4-U).

In the simulations of the closed Lck-FL, the lipid interaction profile of the SH4-U, SH3, and kinase domains remained fairly similar to the simulations of the open Lck-FL. However, in the closed Lck-FL, the SH2 domain exhibited a distinct PIP lipid binding site (referred hereon as secondary PIP binding site) consisting of residues H148, R219, and R222 (**Fig 3C**). Note that K182 and R184 present in the primary binding site interacted with PIP lipids in both open and closed Lck-FL. However, in the closed Lck-FL, PIP interaction was mostly observed in the secondary binding site via residues H148, R219, R222.

### Lck-SH2 adopts a secondary PIP lipid binding site upon mutation of the primary binding site

To investigate the significance of the primary PIP lipid binding site of the open Lck-FL identified above, we mutated its residues i.e., R134A, K135A, K179A, K182A, R184A. This mutation (referred as mut5) in the open Lck-FL led to the loss of PIP interaction via the primary binding site but formed contacts with PIPs via the secondary binding site (**Fig 3D**), thereby resembling the lipid interaction profile of the closed Lck-FL (**Fig 3C**). The lipid interactions and orientations of the other domains remained unaffected by this mutation in the SH2 domain.

The fact that the secondary binding site of Lck-SH2 (H148, R219, R222) is located on the opposite side of the primary binding site (**Fig 3E**) and dominated PIP interaction in the closed Lck-FL suggests that the closed conformation potentially alters the preferred/primary membrane-binding orientation of Lck-SH2 to some degree. This suggests that Lck-SH2 can attain a secondary membrane-bound conformation but is less preferred and potentially weaker. This secondary binding site was observed frequently in the closed state of Lck. It was shown that Lck exhibits lesser membrane binding if its preferred PIP lipid binding site K182/R184 is altered (Sheng et al., 2016). The fact that Lck-membrane binding was reduced and not completely diminished indicates that this secondary PIP lipid binding site may aid membrane association to a certain degree but reduce colocalization with stimulated TCR-CD3 due to change in its orientation. During spatial re-organization of Lck with TCR-CD3 upon activation (Rossy, Williamson, & Gaus, 2012), this alteration of SH2 domain orientation and PIP lipid binding site may also reduce its competence with the preferred open Lck conformation as previously suggested (Hilzenrat et al., 2020; Rossy et al., 2013).

### Simulations of the isolated domains reveal similar interaction with PIP lipids

Following our investigation of lipid interactions of the Lck-FL open and closed states, we also simulated the Lck-SH2, SH3, SH4 domains individually to be able to analyse their lipid interactions independently of the influence of the other domains. Given that the kinase domain is the largest domain, constituting greater than half of the Lck-FL sequence, we also simulated the Lck-SH domains combined (Lck-SH2,3,4-U) to assess their lipid interactions without the influence of the kinase domain. In all the individual Lck-SH and SH2,3,4-U simulations, the protein structure was placed ~6 nm away from the bilayer (**Fig 3A**) to allow it to explore all possible orientations in solution before binding to the membrane.

These simulations suggested that R134, K135, K179, K182, R184 of Lck-SH2 were the most interactive residues with PIP lipids, preferring PIP_3_ over PIP_2_ (**Fig 3F, 3G**) as suggested by previous experimental studies (Sheng et al., 2016) and by our simulations with the Lck-FL models in this study. Lck-SH2 was also found to bind to the membrane within 700 ns of simulation time in all simulations (**Fig S1C**). The SH3 domain required somewhat more simulation time i.e., 1 μs to bind to the membrane. R89 of Lck-SH3 was the most interactive residue with PIP lipids in good agreement with the simulations of the Lck-FL models. Lck-SH4-U made a very small number of contacts when simulated individually (**Fig 3F**) due to the lack of a strong PIP lipid binding site and because the myristoylated and palmitoylated lipid tails failed to insert into the membrane in the majority of the simulations (**Fig S1C**). As a result, we found a significantly larger fraction of SH4-U unbound to the membrane compared to the SH2 and SH3 (**Fig S1D**). Note that, in some individual SH4-U simulations where its lipid tails inserted into the membrane, the SH4-U domain stayed membrane-bound for the rest of the simulation time (**S1 Movie**).

In the Lck-SH2,3,4-U simulations, the Lck-SH2 dominated the interactions with PIP lipids (with R134, K135, K182, R184), while those of SH3 (K84) and SH4-U (R45) were observable but not significant (**Fig 3G**). This indicated that Lck-SH2 lipid interactions were not influenced by the other SH domains. Note that, although the protein had achieved a membrane bound state via the SH2 domain (by ~1.5 μs in all simulations) (**Fig S1C bottom**), the SH4 lipid tails had not inserted in the membrane. The tails were found binding to a small cavity near the SH2-SH3 linker before the protein attained its membrane-bound state (**Fig S1E**) and did not insert into the membrane despite lipid binding initiated by the SH2 domain. This is presumably due to strong hydrophobic interactions between the lipid tails and the SH2-SH3 linker region, and possibly energetically unfavourable to switch to a membrane inserted state. However, it is important to note that membrane insertion may be achieved given more simulation time. Consistent with this observation, *in vitro* studies have reported that the N-terminal myristoyl group in c-Src binds to the SH3 domain while in solution and modulates membrane anchoring (Le Roux et al., 2019).

### Simulations of Lck-FL indicate flexibility of the kinase domain in the open conformation

We deduced the most observed conformations of the membrane bound Lck-FL in its open state from the CGMD simulations using clustering analysis and with a 0.35 nm RMSD cut-off. In the top three most observed conformations of Lck-FL, we observed that Y394 and Y505 often switched positions i.e., in one conformation, Y394 is proximal to the SH2 domain whilst in another conformation, Y505 is proximal to the SH2 domain (**Fig 4A**). This indicates that the kinase domain can rotate and re-orient relative to the SH2 domain. We also performed atomistic MD (ATMD) simulations of the Lck-FL open state in solution (250 ns × 3 replicas) and found similar activity of the kinase domain, where pY394 and Y505 switched positions alternating their proximity to the SH2 domain (**Fig 4B**).

**Fig 4.**
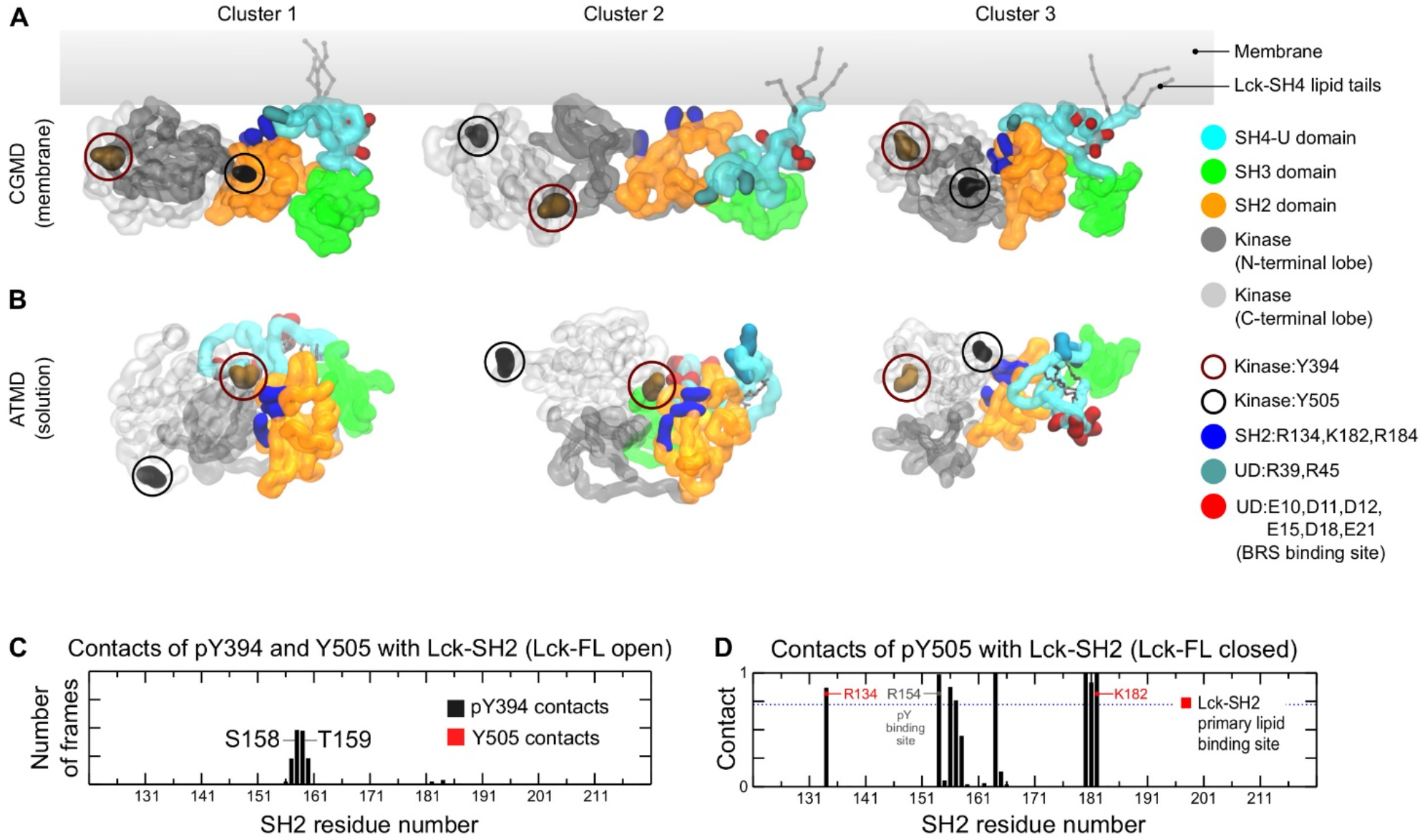
Top three representative conformations of Lck-FL open. **(A)** from CGMD (top) and **(B)** ATMD simulations (bottom) derived from clustering analyses (see main text for details). The kinase domain is made transparent to clarify the positions of Y394 and Y505. Y505 and Y394 are shown in circles to indicate the rotation of the kinase domain relative to the SH2 domain. This is observed by Y505 and Y394 exchanging positions relative to Lck-SH2, in both the membrane associated form of Lck (CGMD) and in solution (ATMD). **(C)** Normalized average number of contacts of pY394 and Y505 with Lck-SH2 in the open Lck-FL state. **(D)** Normalized average number of contacts of pY505 with Lck-SH2 domain in the Lck-FL closed state in ATMD simulations. Normalization was done by dividing the number of contacts by the number of simulation frames thereby obtaining a scale of 0 to 1.

This flexibility of the kinase domain in the Lck-FL open conformation is potentially key to the dynamics of its catalytic activity. Note that, in the ATMD simulations, despite taking up positions near the SH2 domain, neither pY394 nor Y505 contacted the Lck-SH2 PIP lipid binding site (**Fig 4C**) suggesting that Lck-SH2 is free to bind to the membrane in the open state, unlike in the closed state where pY505 interacted with some PIP lipid binding residues of Lck-SH2 (**Fig 4D**).

### Atomistic simulations of the Lck-SH4-U domain reveal its anionic patch

In addition to the open Lck-FL in solution, we performed ATMD simulations of the closed Lck-FL in solution (250 ns × 3 replicas). From these simulations, clustering analyses were performed on the SH4-U domain alone. As a result, we obtained a structure of the most observed conformations of the Lck-SH4-U in the open and closed states of Lck. We then calculated their electrostatic profiles and compared them with the electrostatic profile of the initial SH4-U model obtained earlier in this study (**Fig 5**). This revealed that the SH4-U domain, despite its dynamic nature, maintained an anionic patch (E10, D11, D12, E15, D18, E21) which was independent of the open and closed Lck-FL conformations. Interestingly, this anionic patch was identified on the opposite side of its PIP lipid binding surface (R39, R45) indicating its availability to bind to cationic residues of other proteins, especially those of the BRS motifs of CD3ε and ζ subunits of the TCR-CD3 complex as suggested by NMR experiments (Li et al., 2017).

**Fig 5.**
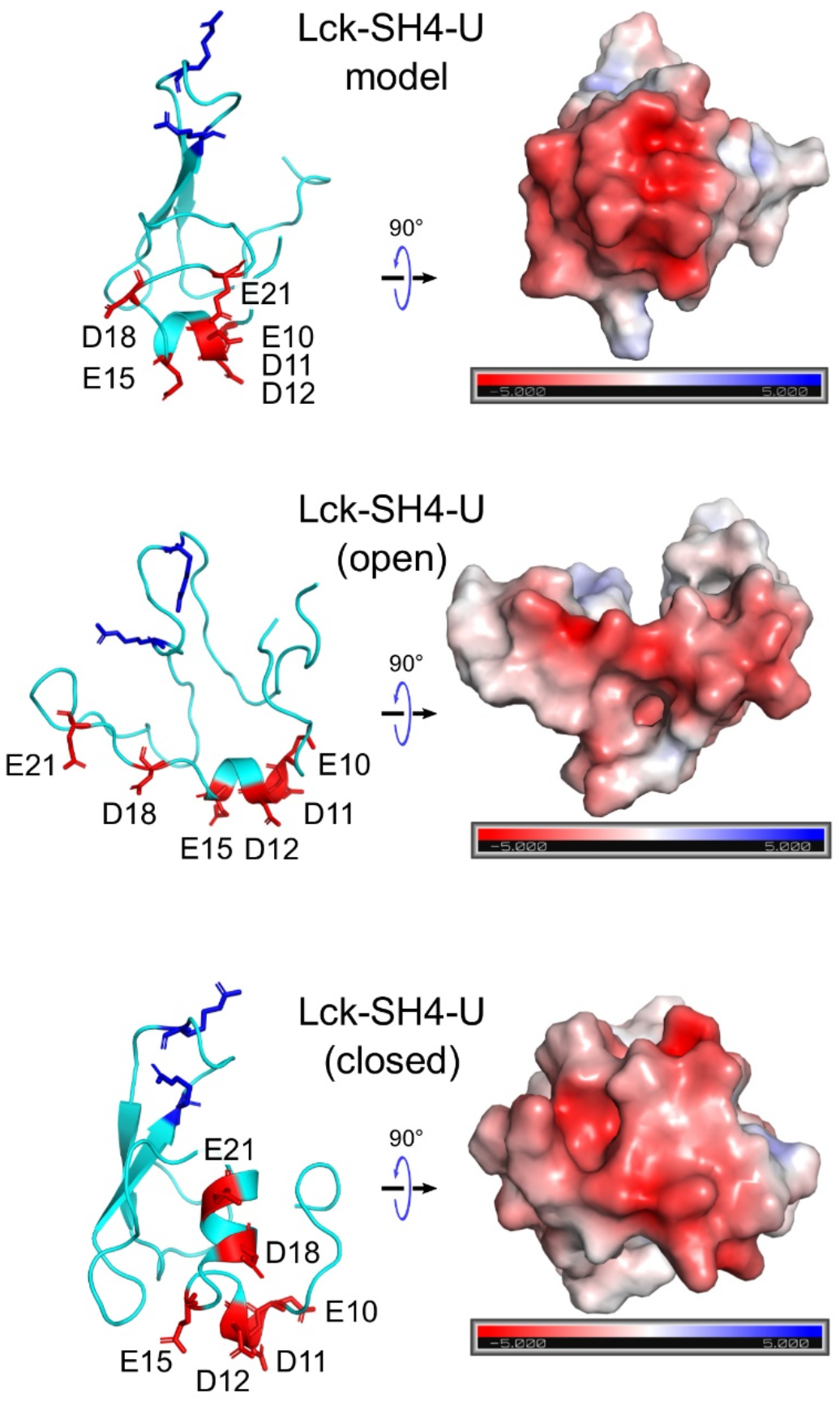
***The electrostatic profiles of the Lck-SH4-U domain** as in the initial model (top), and according to clustering analyses after simulating the Lck-FL open state (middle), and the Lck-FL closed state (bottom). Their electrostatic profiles were calculated in the ±5 kT/e range and at pH 7.0 using the PDB2PQR* (Dolinsky et al., 2004) *and APBS* (Baker et al., 2001) *tools. Electronegative and electropositive regions are indicated by the red and blue intensities respectively. The residues forming the anionic patch are shown as red sticks and labelled. PIP lipid binding residues (R39, R45) are shown as blue sticks for reference*.

## CONCLUSION

In this study, we have revealed lipid interactions of the full-length post-translationally modified Lck in both its open and closed conformations and highlighted its PIP lipid binding sites. Our key finding concerning PIP lipid binding was that the SH2 domain adopts a secondary binding site in the closed state of Lck-FL compared to its open state. Although this secondary binding site may aid in membrane localization to some degree, it may be less important during spatial organization of open Lck during T cell activation. Our simulations show that upon membrane binding of Lck, it is surrounded by a pool of negatively charged lipid headgroups creating an anionic environment, whereas it is also observed that the TCR-CD3 cytoplasmic tails maintain an anionic lipid environment (Prakaash et al., 2021). This sheds light on the potential significance of lipids during TCR-Lck association.

In this study, we suggest that the residues R134, R135, and K179 contribute to the primary PIP lipid binding site in addition to those previously reported i.e., K182 and R184 (Sheng et al., 2016). Moreover, all five of these residues are found to be conserved among Src family members (**Fig S2A**) indicating that all their SH2 domains are likely to localize to the membrane with the same orientation. The PIP interactions of Lck-SH3, although not significant, consistently interacted via K84. Upon lipid tail insertion into the membrane, the Lck-UD also showed significant interaction with PIPs via R39 and R45. Furthermore, we present a structural model of the Lck-UD that reveals an anionic patch with which it could bind to basic-rich motifs of the TCR-CD3 subunits and aid in TCR-Lck association.

We also found that the kinase domain interacts with PIPs in the membrane via residues R455, R458, R474, K478 which form a cationic patch at the bottom of its C-terminal lobe. These residues, along with other potentially PIP contacting residues, were also found to be conserved among other Src family members (**Fig S2A**) suggesting similar lipid interactions and membrane-bound orientations of the kinase domains in the Src family. Further, the lipid interactions of the kinase domain implies that it is likely to be situated proximal to the membrane surface. Therefore, given that the TCR-CD3 cytoplasmic region is also closely associated with the membrane surface (Prakaash et al., 2021), it is likely that Lck kinase-mediated phosphorylation of TCR-CD3 ITAMs and of other downstream signalling proteins during the initial phase of T cell activation is carried out proximal to the plasma membrane. Similarly, given the conservation of important PIP binding sites in the SH2 and kinase domains, ITAM phosphorylation mediated by other members of the Src family may also occur close to the surface of plasma membrane.

It is also important to consider some limitations of this study. Here, we performed CGMD simulations using the MARTINI forcefield (de Jong et al., 2013) which involved elastic network (EN) restraints (Periole, Cavalli, Marrink, & Ceruso, 2009) within each domain of the open state of Lck in order to maintain their tertiary structures as suggested by experiments (PDB:4D8K, 3LCK). To make the open Lck simulations more realistic, we avoided inter-domain restraints allowing each domain to freely associate with each other. In the closed state of Lck, it is known that the kinase binds to the SH2 and SH3 domains. Therefore, such a configuration was homology modelled based on the structure of the closed state of Hck, a member of the Src kinase family (PDB:5H0B) (Yuki et al., 2017) found to exhibit highest identity with Lck among other Src members (**Fig S2B**). This homology model of Lck was restrained using EN in CGMD simulations to retain its closed conformation. The SH4-U domain was modelled based on secondary structure predictions, validated using ATMD simulations and available experimental evidence, and finally conjoined with the rest of the Lck structure in both open and closed conformations.

## METHODS

### Molecular Modelling

To obtain a model of the 3D structure of Lck-SH4-U, the PSIPRED secondary structure prediction tool (Buchan & Jones, 2019) along with 3D structure predictions by the I-Tasser (Yang et al., 2015) and Robetta (D. E. Kim et al., 2004) servers were used. The sequence of the SH4-U domain was obtained from UniprotKB (P06239). Post-translational acylations/lipid tails were added using CHARMM-GUI (Jo, Kim, Iyer, & Im, 2008). Modeller 9.2 (Eswar et al., 2006; Webb & Sali, 2014) and UCSF Chimera (Pettersen et al., 2004) were used to conduct modelling of the open Lck-FL. Homology modelling of the closed Lck-FL was conducted based on available structural data of Hck using Modeller 9.2.

### Coarse-grained molecular dynamics (CGMD) simulations

All models were coarse-grained using the Martini 2.2 forcefield (Marrink, Risselada, Yefimov, Tieleman, & de Vries, 2007) and the *martinize* script. To coarse-grain the lipid tails along with the rest of the protein, the *martinize* script, and the Martini 2.2 amino acid topology were modified to include published parameters (Atsmon-Raz & Tieleman, 2017), and made publicly available (https://github.com/DJ004/martini_mod).

CGMD simulations were set up using the *Insane* tool (Wassenaar, Ingólfsson, Böckmann, Tieleman, & Marrink, 2015) and Gromacs 5.0. EN restraints (Periole et al., 2009) with a 1000 kJ/mol/nm^2^ force constant and 0 to 0.7 nm cut-off distance was applied. However, the restraints were applied only within each domain to maintain their tertiary structure and not between domains to allow unbiased inter-domain interactions. Membrane lipid compositions used to set up each CGMD simulation are shown in Table 1. In all CGMD simulations, each lipid contained one saturated acyl chain and one mono-unsaturated acyl chain, while their headgroup composition is based on the composition of TCR-CD3 activation domains in the T cell plasma membrane (Zech et al., 2009). The solvent was neutralized with 0.15M Na+ and Cl- ions. All systems were energy minimized using the steepest descent algorithm until the maximum force converged to 1000 kJ/mol/nm and equilibrated for 2.5 ns with the protein position-restrained. The equilibrated system was then used to generate differing initial velocities for twenty production simulations run for 5 μs each with a 20 fs time-step. The NPT ensemble was used to conduct equilibration and production simulations. Co-ordinates were saved at 200 ps intervals. A semi-isotropic Parrinello-Rahman barostat (1 bar) (Parrinello & Rahman, 1981) and V-rescale thermostat (323 K) (Bussi, Donadio, & Parrinello, 2007) were used for production simulations along with a 3×10^−4^/bar compressibility.

### Atomistic molecular dynamics (ATMD) simulations

CHARMM-GUI (Jo et al., 2008) was used with the CHARMM36 forcefield (Huang & MacKerell, 2013) to setup ATMD simulations of the initial Lck-SH4-U model, the Lck-FL open model and closed model in solution using the TIP3 water as solvent neutralized with 0.15M Na+ and Cl- ions. All systems were energy minimized using the steepest descent algorithm using Gromacs 2016 until the maximum force converged to 1000 kJ/mol/nm^2^, followed by isotropic (NPT) equilibration at 323 K where the protein backbone was position-restrained. The equilibrated system was used to generate differing initial velocities for three production simulations run 250 ns each using a 2 fs time-step. Co-ordinates were saved at 40 ps intervals. The V-rescale thermostat (323 K) (Bussi et al., 2007) and Parrinello-Rahman isotropic barostat (1 bar) (Parrinello & Rahman, 1981) was used with a compressibility of 4.5×10^−5^/bar. The LINCS algorithm (Hess, Bekker, Berendsen, & Fraaije, 1997) applied constraints on hydrogen bond lengths and the Particle Mesh Ewald algorithm (Essmann et al., 1995). Coulombic and van der Waals interactions were defined by a 1.2 nm distance cut-off.

### Data analysis and visualization

Protein-lipid and protein-protein interactions in CGMD simulations were calculated using the *gmx mindist* command where a contact was defined by a 0.55 nm distance cut-off. All contact analyses results represent merged data from all simulation replicates. Clustering analyses used the *gmx cluster* command and the *gromos* method (Daura et al., 1999) with an RMSD cut-off of 0.35 nm. For this, all trajectories were concatenated using gmx *trjcat*, the protein was extracted using *gmx trjconv* and RMSD calculations were run skipping 5 frames for both CGMD and ATMD. Distance versus time and radial distribution function calculations were done using the *gmx distance* and *gmx rdf* commands respectively. VMD was used for visualization and rendering (Humphrey, Dalke, & Schulten, 1996). The APBS (Baker et al., 2001) plugin of PyMOL 2.4 (pymol.org) was used to calculate electrostatics. Xmgrace (https://plasma-gate.weizmann.ac.il/Grace/) and Matplotlib 3.3 (doi.org/10.5281/zenodo.3948793) were used for plotting.

## ACKNOWLEDGMENTS

This research was supported by ARC3 and ARC4 supercomputers, part of the High Performance Computing facility at the University of Leeds, Leeds, United Kingdom. O.A. is funded by the Wellcome Trust grant 200844/Z/16/Z. For the purpose of open access, the author has applied a CC BY public copyright licence to any Author Accepted Manuscript version arising from this submission.

## DATA AVAILABILITY

Simulation data will be available via https://doi.org/10.5518/1158

## SUPPORTING INFORMATION

**Fig S1.**
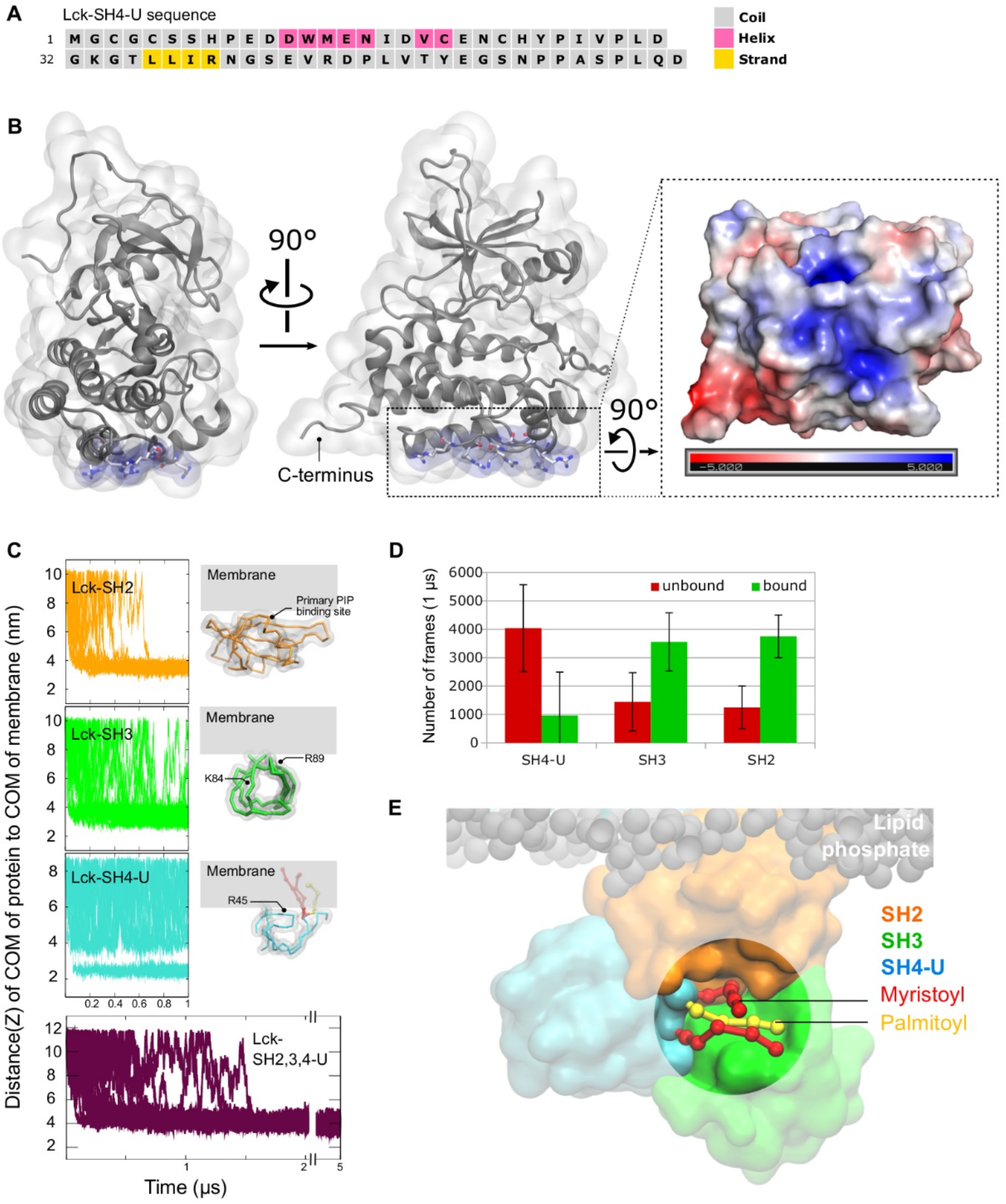
Secondary structure prediction of Lck-SH4-U, membrane binding of individual Lck-SH domains, and electrostatics of the kinase domain. **(A)** Secondary structure prediction of Lck-SH4-U by PSIPRED 4.0 server. **(B)** Residues in the kinase domain forming a flat cationic patch. Their electrostatic profile was calculated using in the ±5 kT/e range and at pH 7.0 using the PDB2PQR and APBS tools. Electronegative and electropositive regions are indicated by red and blue intensities respectively. **(C)** Distance between the center of mass (COM) of protein to COM of membrane in all 20 simulation replicates: SH4-U (cyan), SH3 (green), SH2 (orange), Lck-SH2,3,4-U (maroon). The most observed membrane-bound orientations of the individually simulated Lck-SH domains are also shown. Residues interacting most with the membrane in these simulations are labelled. **(D)** Average number of frames in the individual Lck-SH simulations that the protein stayed unbound (red) or bound (green) to the membrane. **(E)** A snapshot from one of the Lck-SH2,3,4-U simulations highlighting the binding pocket of the SH4 lipid tails (near the SH2-SH3 linker loop region) when they did not insert into the membrane.

**Fig S2.**
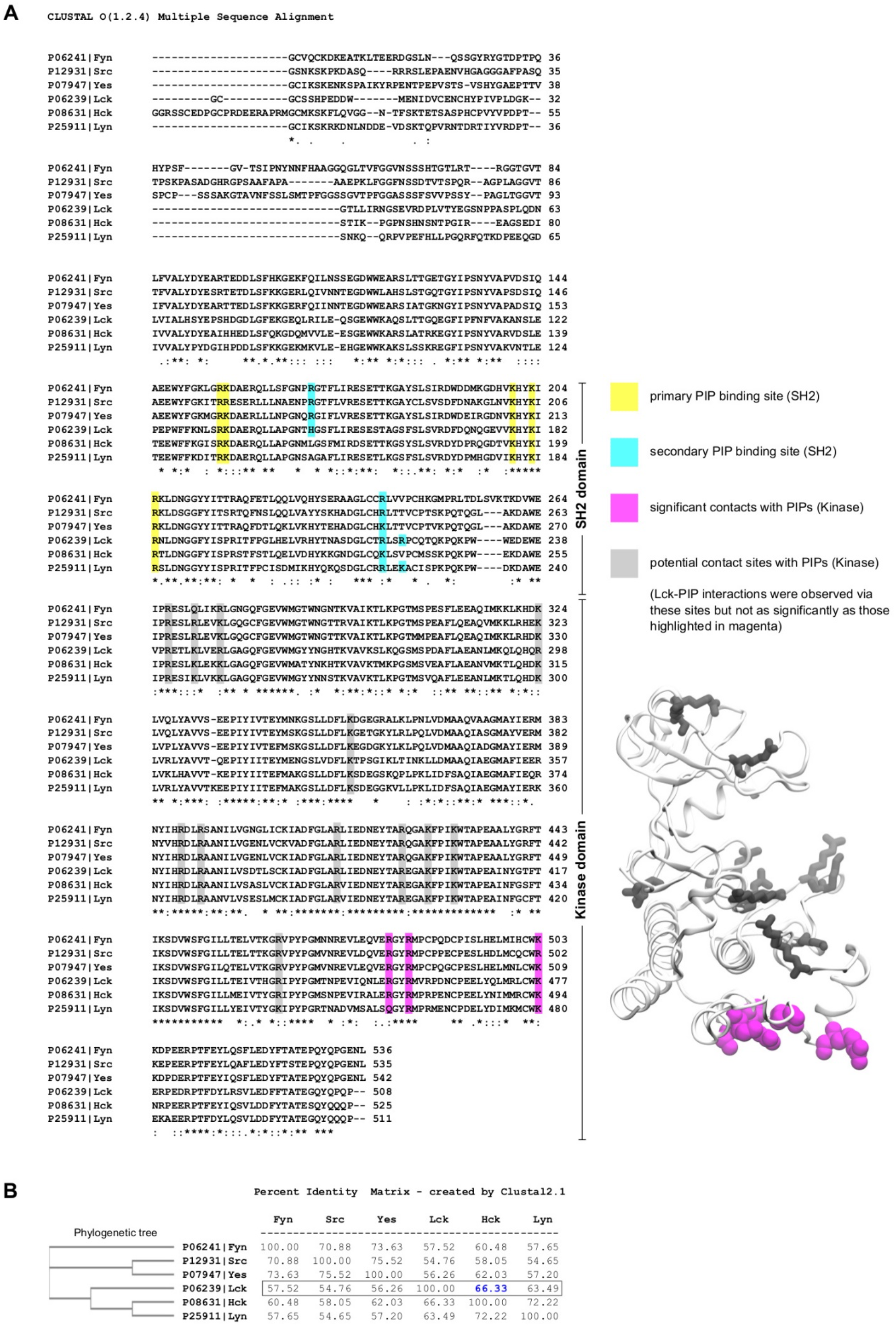
Multiple sequence alignment and the identity of the full-length Lck with other members of the Src family of kinases. **(A)** Multiple sequence alignment, obtained by Clustal Omega, highlighting the residues in the SH2 and kinase domains that mediated Lck-PIP interactions. These residues are observed to be conserved. Note that the myristoylated G2 residue is considered as the first residue, hence all residues are shifted one position behind i.e., ‘n-1’. Refer **Fig 3E** to visualize the primary and secondary PIP lipid binding sites of Lck-SH2. The figure shown on the right is a cartoon representation of the kinase domain with significant PIP contacting residues shown as magenta speheres, and potentially contacting residues shown as grey sticks. **(B)** Schematic phylogenetic tree showing the evolution of Src family of kinases and their identity matrix (calculated in %). This suggests that Lck is most similar to Hck compared to other Src family members. The protein sequences were obtained from Uniprot (whose IDs are listed beside the respective protein names).

**S1 Movie. CGMD simulation displaying insertion of myristoylated and palmitoylated lipid tails of the SH4 domain into the membrane.** Phospholipid tails are shown as transparent grey spheres and their headgroups as transparent coloured spheres (POPC: brown, POPE: purple, PIP_2_: orange, PIP_3_: red). The myristoylated and palmitoylated residues are shown as yellow and red ball and sticks respectively. Lck-SH4-U backbone is shown as white bonds, and its PIP lipid binding residues R39, R45 are shown as blue surfaces.

